# A modular system to label endogenous presynaptic proteins using split fluorophores in *C. elegans*

**DOI:** 10.1101/2024.07.29.605690

**Authors:** Mizuki Kurashina, Andrew W. Snow, Kota Mizumoto

## Abstract

Visualizing the subcellular localization of presynaptic proteins with fluorescent proteins is a powerful tool to dissect the genetic and molecular mechanisms underlying synapse formation and patterning in live animals. Here, we utilize split green and red fluorescent proteins to visualize the localization of endogenously expressed presynaptic proteins at a single neuron resolution in *Caenorhabditis elegans.* By using CRISPR/Cas9 genome editing, we generated a collection of *C. elegans* strains in which endogenously expressed presynaptic proteins (RAB-3/Rab3, CLA-1/Piccolo, SYD-2/Liprin-α, UNC-10/RIM and ELKS-1/ELKS) are tagged with tandem repeats of GFP_11_ and/or wrmScarlet_11_. We show that the expression of wrmScarlet_1-10_ and GFP_1-10_ under neuron-specific promoters can robustly label presynaptic proteins in different neuron types. We believe that combinations of knock-in strains and wrmScarlet_1-10_ and GFP_1-10_ plasmids are a versatile modular system to examine the localization of endogenous presynaptic proteins in any neuron type.

## Introduction

The synapse is the basic functional unit of the nervous system and consists of the presynaptic specialization that sends a chemical signal via exocytosis of neurotransmitter-containing synaptic vesicles (SVs) and the postsynaptic specialization that receives neurotransmitters through postsynaptic receptors (Figure 1A). The presynaptic specialization is defined by the cluster of SVs and the electron-dense region known as the active zone^1,2^, where SNARE-complex dependent SV exocytosis occurs^3^. The active zone consists of a group of conserved proteins known as active zone proteins which includes Liprin-α, ELKS, RIM (Rab3-interacting molecule), RIM-BP (RIM-binding protein), Munc13 (Mammalian UNC-13), and Piccolo/Bassoon. The functional and structural homologs in *Caenorhabditis elegans* also play crucial roles in the formation and function of the active zone, and these homologs include SYD-2/Liprin-α, ELKS-1/ELKS, UNC-10/RIM, RIMB-1/RIM-BP, UNC-13/Munc13, and CLA-1/Piccolo/Bassoon. The functions of these active zone proteins are summarized in a recent review^4^.

**Figure 1.**
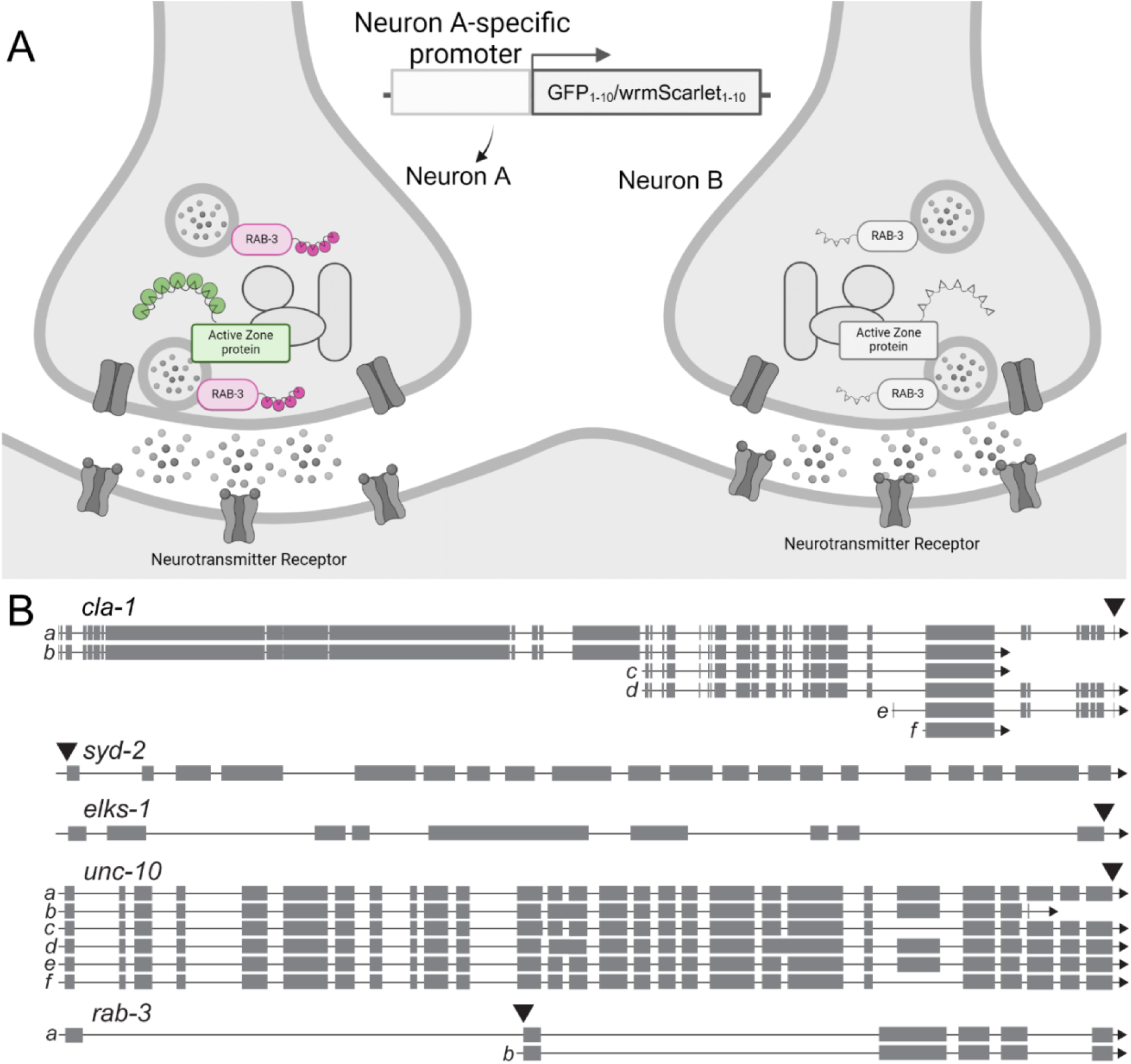
Schematic of split fluorescent protein system and insertion sites. (A) Schematic of the presynaptic structure of two neurons (Neuron A and B) and the split fluorescent protein system to label presynaptic proteins in a specific neuron. An active zone protein and RAB-3 are tagged with seven tandem repeats of GFP_11_ and four tandem repeats of wrmScarlet_11_, respectively (represented by triangles). Neuron A-specific expression of GFP_1-10_ and wrmScarlet_1-10_ specifically reconstitutes fluorescent GFP and wrmScarlet in neuron A but not in neuron B. Created with BioRender.com. (B) Schematics of the genomic regions of *cla-1, syd-2, elks-1, unc-10,* and *rab-3.* The knock-in sites of *GFP_11_* and/or *wrmScarlet_11_* are indicated by the black arrowheads.

In *C. elegans*, transgenic expression of fluorescently tagged presynaptic proteins under neuron-type specific promoters has been widely used to visualize presynaptic structures in live animals^4,5^. Particularly, forward genetic screenings using the transgenic animals expressing fluorescently tagged SV-associated proteins (SNB-1/Synaptobrevin and RAB-3/Rab3) have played key roles in identifying genes essential for presynaptic assembly and specificity^6–8^. On the other hand, labeling active zone proteins by using traditional transgenic overexpression of fluorescently-tagged active zone proteins is more challenging, as the overexpression of these proteins often results in aberrant subcellular localization^9,10^ that does not recapitulate the active zone structure observed by electron microscopy^9,11,12^. This is likely because overexpression of active zone proteins affects proper presynaptic assembly. Visualizing endogenous active zone proteins by knocking-in fluorescent protein sequences using CRISPR/Cas9 mitigates the issues associated with transgenic overexpression. However, this method labels in all the cells in which the tagged protein is expressed and therefore cannot be used to examine the changes to protein subcellular localization in a single neuron^13^. It is also possible that the expression level of the endogenous protein may be too low to visualize when a single fluorescent protein is fused. The recent employment of split-fluorescent proteins to visualize endogenous proteins can mitigate both limitations described above. By tagging proteins of interest with tandem repeats of one fragment of the fluorescent protein and expressing the remaining part of the fluorescent protein under a tissue-specific promoter, an endogenously expressed protein can be labeled with multiple copies of fluorescent proteins in a cell-type specific manner^14–18^.

Here by using CRISPR/Cas9 genome editing, we generated a collection of *C. elegans* strains to visualize endogenous presynaptic proteins in a neuron-specific manner. We knocked-in sequences of tandem repeats of the last β-sheet of GFP (GFP_11_)^14–17^ and/or wrmScarlet (wrmScarlet_11_)^18^ into the genomic loci of *rab-3* and active zone genes (*syd-2, elks-1, unc-10,* and *cla-1*), and show that these strains can visualize endogenous presynaptic proteins in neuron-specific manners when the remainder of the fluorescent protein (GFP_1-10_ and wrmScarlet_1-10_) are expressed under the neuron type-specific promoters. We believe that these strains are useful to researchers to examine the presynaptic specialization in the neuron-types of their interest.

## Materials and Methods

### C. elegans strains

Bristol N2 strain was used as a wild-type reference. All strains were cultured in the nematode growth medium (NGM) with OP50 as described previously^19^. All strains were maintained at room temperature (22°C). The following alleles were used in this study: *rab-3(miz237[4×wrmScarlet_11_::rab-3])* II*, cla-1(miz321[cla-1::7×GFP_11_])* IV*, cla-1(miz329[cla-1:: 8×wrmScarlet_11_])* IV, *elks-1(miz364[elks-1::7×GFP_11_])* IV*, unc-10(miz404[unc-10::7×GFP_11_])* X*, syd-2(miz231[7×GFP_11_::syd-2])* X*, syd-2(miz329[8×wrmScarlet_11_::syd-2])* X*, syd-2(ok217) X*.

### Transgenes

The following transgenes were used in this study:

*mizIs41*(*mig-13*p::*wrmScarlet_1-10_::SL2::GFP_1-10_* (10 ng/μL)*; odr-1*p*::GFP*), *mizEx605*(*itr-1*p::*wrmScarlet_1-10_::SL2::GFP_1-10_* (20 ng/μL)*; odr-1*p*::GFP*), *mizEx657*(*itr-1*p::*wrmScarlet_1-10_::SL2::GFP_1-10_* (200 ng/μL)*; odr-1*p*::GFP*), *mizEx607*(*dat-1*p*::wrmScarlet_1-10_::SL2::GFP_1-10_* (10 ng/μL*); odr-1*p*::GFP*), *mizEx624(unc-25*p*::wrmScarlet_1-10_::SL2::GFP_1-10_* (2 ng/μL*); odr-1*p*::GFP*), *wyIs685*(*mig-13*p::*TdTomato::rab-3b, mig-13*p*::3×GFPnovo2::cla-1; odr-1*p*::GFP*). The *mizIs41* transgene was obtained by a spontaneous integration when trying to generate the extrachromosomal arrays for *mig-13*p::*wrmScarlet_1-10_::SL2::GFP_1-10_*.

The transgenic lines with extrachromosomal arrays were generated using the standard microinjection method^20,21^. The *odr-1*p*::GFP* co-injection marker plasmid was injected at 20 ng/μL.

### Plasmid construction

*C. elegans* expression plasmids were made in a derivative of pPD49.26 (A. Fire), the pSM vector. To visualize the presynaptic specializations of the DA9 neuron, VD and DD neurons and the PDE neurons, we cloned the DA9 specific promoters (*itr-1B*p^22^ or *mig-13*p^23^)^24^, VD/DD-specific promoter (*unc-25*p)^25^, PDE-specific promoter(*dat-1*p)^10,26^ into the *Sph*I and *Asc*I sites of the *wrmScarlet_1-10::_SL2::GFP_1-10_* plasmid.

*4×wrmScarlet_11,_ 7×GFP_11_, 8×wrmScarlet_11_* sequences were synthesized using GeneArt Gene Synthesis service (ThermoFisher Scientific) and were cloned into either their default plasmids or pBluescriptII (SK-). These plasmids were used as a PCR template to amplify the homology directed repair (HDR) template. The sequences of the HDR templates and primers used to amplify the HDR templates are available in the supplemental information.

### CRISPR/Cas9 genome editing

Insertion of the tandem repeats of *GFP_11_* and/or *wrmScarlet_11_* into the loci of presynaptic genes was conducted using CRISPR/Cas9 genome editing according to the protocols described previously^27,28^. We amplified the donor HDR template of the tandem repeats of *GFP_11_* and/or *wrmScarlet_11_* with 50-60 bp of homology arm sequences to each of the synaptic gene loci using Phusion or Q5 high-fidelity DNA polymerases (New England Biolabs). The PCR products were ‘melted’ and injected along with the Cas9-gRNA RNP complex and a pRf4 *rol-6* co-injection marker plasmid^21^ into both gonad arms of gravid worms. F1s were singled and screened for knock-in by PCR genotyping. The insertions and their flanking sequences were confirmed by Sanger sequencing. The primer sequences for amplifying the HDR templates and genotyping are listed in the supplemental information.

### Aldicarb assay

The stock solution of aldicarb (Sigma-Aldrich: 33386-100MG), prepared at 100 mM in 70% ethanol, was added to the NGM to a final concentration of 1 mM concentration after autoclaving and cooling to 55 °C. 25-30 adult animals were transferred to the 1mM aldicarb plates and scored for paralysis every 20 minutes. We defined animals as paralyzed when they were completely motionless and unresponsive when prodded with a platinum wire 3 times. Each genotype was assayed 3 times.

### Confocal Microscopy

Images of fluorescently tagged fusion proteins were captured in live *C. elegans* using a Zeiss LSM800 Airyscan confocal microscope (Carl Zeiss, Germany) equipped with a 63× magnification oil objective lens (Carl Zeiss, Germany). Worms were immobilized using a 3:1 mixture of 0.225 M 2,3-butanedione monoxime (Sigma-Aldrich) and 7.5 mM levamisole hydrochloride (Sigma-Aldrich) and mounted on 2.5% agarose pads. Images were analyzed and processed using Zen software (Carl Zeiss) and ImageJ (NIH, USA).

### Fluorescent signal intensity quantification

Signal intensity of CLA-1::7×GFP from 35 z-stacks (0.15 μm thickness) of ∼50 μm segment of the DA9 synaptic domain were examined in N2 animals. Using ImageJ, images were processed via Z-projection sum slices and straightened. Particle thresholding and analysis were conducted by first subtracting background fluorescence and applying Gaussian blur. Auto-local thresholding using Bernsen’s thresholding was applied using the default values. Regions of interest (ROI) denoting the CLA-1 puncta were manually examined to confirm visible fluorescence. Particles were then measured to obtain the mean fluorescence value of each ROI.

### Statistics

Prism10 (GraphPad Software, USA) was used for statistical analyses. We applied the one-way ANOVA method with post hoc Tukey’s multiple comparison test for comparison among three or more parallel groups with multiple plotting points, and log-rank survival test with Bonferroni correction for comparison of paralysis on aldicarb. Data were plotted with error bars representing standard deviation (SD).

## Results

### Generating GFP_11_ and wrmScarlet_11_ knock-in strains to visualize endogenous presynaptic proteins

We employed split-GFP and split-wrmScarlet to label endogenous presynaptic proteins in a neuron-specific manner^15–18^. Split-wrmScarlet is an engineered split fluorophore that has been codon-optimized for *C. elegans* and is a derivative of yeast codon-optimized mScarlet^18^. Specifically, we knocked-in seven tandem repeats of *GFP_11_* (*7×GFP_11_*) or eight tandem repeats of *wrmScarlet_11_* (*8×wrmScarlet_11_*) sequences to the core active zone genes (*syd-2, elks-1, unc-10,* and *cla-1*) using the CRISPR/Cas9 genome editing technology (Figure 1B). *7×GFP_11_* and *8×wrmScarlet_11_* were inserted at the 5’ end of *syd-2* locus and at the 3’ end of *cla-1*, respectively. *7×GFP_11_* was inserted at the 3’ ends of *unc-10*, and *elks-1* loci, according to previous works which used transgenic labeling of these active zone proteins^13,29–31^ (Figure 1B). To visualize SVs, four tandem repeats of the *wrmScarlet_11_* (*4×wrmScarlet_11_*) sequence were inserted at the 5’ end of *rab-3 B isoform* (Figure 1B)^13,32–35^.

The fusion of any small sequences including *7×GFP_11_* and/or *8×wrmScarlet_11_* to the endogenous active zone proteins may interfere with their functions as they may cause steric hinderance to other proteins or misfolding. To rule out the potential deleterious effects of the insertion of these sequences into our genes of interest, we first examined the locomotion pattern of the knocked-in strains. The null mutants of *syd-2* and *unc-10* exhibit uncoordinated locomotion (*Unc*) phenotypes^7,19^, while all the knock-in strains we generated, including *7×GFP_11_::syd-2* and *unc-10::7×GFP_11_* are superficially wild-type in terms of their locomotion (Figure S1 and not shown). This suggests that the *7×GFP_11_* and *8×wrmScarlet_11_* tags do not abolish the functions of these proteins.

While the locomotion of these animals indicate that they are superficially wild type, we further characterized the effects of GFP_11_ and/or wrmScarlet_11_ knock-ins on neurotransmission by aldicarb assay^36,37^. Previous works showed that null mutants of *cla-1* and *syd-2* are resistant to aldicarb due to impaired neurotransmission^7,13^. We found that that *cla-1::7×GFP_11_* and *7×GFP_11_::syd-2* mutants exhibited normal sensitivity to aldicarb (Figure 2), suggesting that CLA-1::7×GFP_11_ and 7×GFP_11_::SYD-2 fusion proteins retain wild type functions in neurotransmission. Loss of function mutants of *unc-10* also exhibit strong aldicarb resistance^38^. In *unc-10::7×GFP_11_* animals, we observed modest aldicarb resistance (Figure 2), suggesting that the *7×GFP_11_* insertion weakly affects *unc-10* function. Null mutants of *elks-1* do not exhibit any defects in aldicarb sensitivity^39^. Interestingly, we observed that *elks-1::7×GFP_11_* animals exhibit a modest resistance to aldicarb (Figure 2). It is possible that ELKS-1::7×GFP_11_ fusion protein weakly interferes with the functions of other active zone proteins that interact with ELKS-1, thereby affecting neurotransmission. *4×wrmScarlet_11_::rab-3* animals exhibited a strong aldicarb resistance (Figure 2), similar to the null mutant of *rab-3*, as previously described^40,41^. This suggests that 4×wrmScarlet_11_::RAB-3 is not functional in neurotransmission. Nevertheless, the localization patterns of 4×wrmScarlet::RAB-3 is similar to transgenically expressed RAB-3 and other SV markers^7,10,13,42,43^ (see below) (Figures 3E, 3F, 4). While the function of RAB-3 is affected in *4×wrmScarlet_11_::rab-3* animals, we conclude that *4×wrmScarlet_11_::rab-3b* can be used as a SV marker strain.

**Figure 2.**
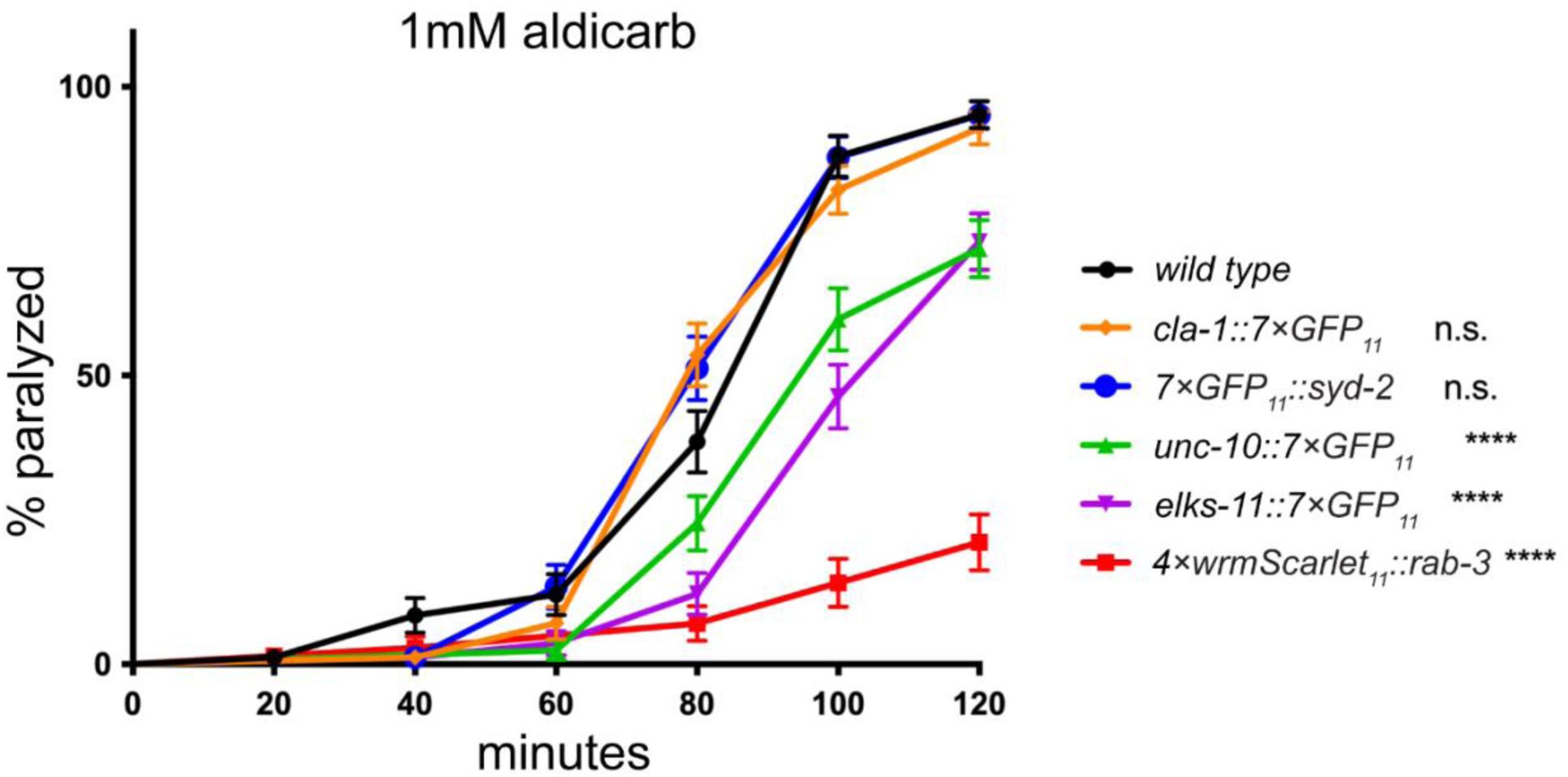
Effects of split-fluorophore sequence knock-ins on neurotransmission. The time course of paralysis for wild type, *cla-1::7×GFP_11_, 7×GFP_11_::syd-2, unc-10::7×GFP_11_*, *4×wrmScarlet_11_::rab-3*, and *elks-1::7×GFP_11_* animals on 1.0 mM aldicarb plates from three independent trials with n > 20 for each genotype in each trial. Log-rank survival test with Bonferroni correction compared to wild type. n.s. not significant; **** p < 0.0001

**Figure 3.**
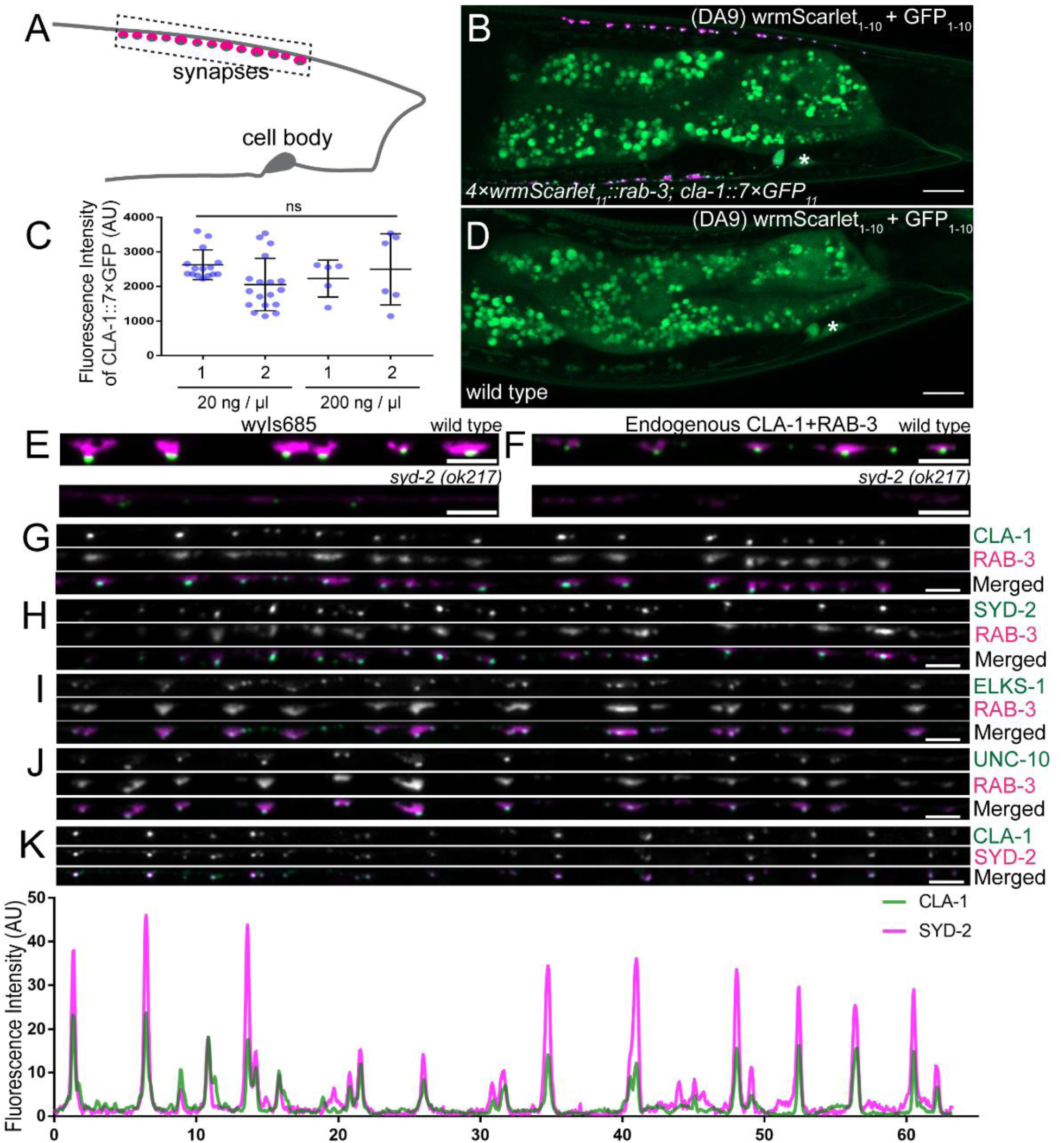
Labeling endogenous presynaptic proteins in DA9. (A) Schematic of the DA9 motor neuron. The dotted box represents the imaging area of the presynaptic domains shown in (G-K). (B) Representative image of the DA9 motor neuron labeled with endogenous 4×wrmScarlet_11_::RAB-3 and CLA-1::7×GFP_11_. The *mig-13* promoter was used to express GFP_1-10_ and wrmScarlet_1-10_. The asterisk denotes the DA9 cell body. Scale bar: 10 μm. (C) Quantification of fluorescence intensity of CLA-1::7×GFP in animals expressing GFP_1-10_ and wrmScarlet_1-10_ under the *itr-1* promoter. The *itr-1*p*::wrmScarlet_1-10_::SL2::GFP_1-10_* plasmids were injected at 20 ng/μL and 200 ng/μL, and two independent lines per each injection were quantified. Each dot represents average CLA-1::7×GFP_11_ signal intensity per synapse per animal. n.s. not significant. (D) Representative image of the DA9 motor neuron expressing GFP_1-10_ and wrmScarlet_1-10_ under the *mig-13* promoter in wild type animals. We observed a dim GFP_1-10_ signal in the cell body of DA9. The asterisk denotes the DA9 cell body. Scale bar: 10 μm. (E) Representative images of a few DA9 presynaptic specializations within the DA9 synaptic domain visualized by the *wyIs685* transgene expressing CLA-1::3×GFPnovo2 (green) and TdTomato::RAB-3 (magenta) in wild type (top) and *syd-2(ok217)* mutant (bottom). Scale bar: 2 μm (F) Representative images of a few DA9 presynaptic specializations within the DA9 synaptic domain visualized by the endogenous CLA-1::7×GFP_11_ (green) and 4×wrmScarlet::RAB-3_11_ (magenta) in wild type (top) and *syd-2(ok217)* mutant (bottom). Scale bar: 2 μm. (G-J) Representative images of the DA9 presynaptic specializations labeled with CLA-1::7×GFP_11_ (top), 4×wrmScarlet::RAB-3 (middle), and merged image (bottom) (G), 7×GFP_11_::SYD-2 (top), 4×wrmScarlet_11_::RAB-3 (middle), and merged image (bottom) (H), ELKS-1::7×GFP_11_ (top), 4×wrmScarlet_11_::RAB-3 (middle), and merged image (I), UNC-10::7×GFP_11_ (top), 4×wrmScarlet_11_::RAB-3 (middle), and merged image (bottom) (J). Scale bar: 2 μm. (K) Representative image of CLA-1::7×GFP_11_ and 8×wrmScarlet_11_::SYD-2 localization in the DA9 neuron. Scale bar: 2 μm. The corresponding fluorescent intensity plots for CLA-1::7×GFP_11_ and 8×wrmScarlet_11_::SYD-2 are shown below.

### Labeling endogenous presynaptic proteins in the DA9 motor neuron

We tested if these knock-in strains can be used to visualize endogenous presynaptic proteins in a neuron-specific manner using the DA9 motor neuron. DA9 is one of nine dorsal A-type (DA) cholinergic motor neurons required for the backward locomotion of *C. elegans*^44^. The cell body of DA9 resides in the ventral side of the worm near the preanal ganglion and sends a dendrite ventrally and an axon dorsally where it forms approximately 20 *en passant* synapses onto the dorsal body wall muscles^45^ (Figures 3A and 3B). Previous works have shown that transgenic overexpression of CLA-1::3×GFPnovo2 and TdTomato::RAB-3 can label the presynaptic structures of DA9; CLA-1::3×GFPnovo2 is localized at the tip of each presynaptic varicosity labeled with TdTomato::RAB-3^13,32^ (Figures 3A and 3I).

We examined the localization patterns of CLA-1::7×GFP_11_ and 4×wrmScarlet_11_::RAB-3 in DA9 using a transgene *mizIs41 (mig-13*p*::wrmScarlet_1-10_::SL2::GFP_1-10_)* expressing wrmScarlet_1-10_ and GFP_1-10_ in DA9 of *4×wrmScarlet_11_::rab-3; cla-1::7×GFP_11_* animals. The localization patterns of 4×wrmScarlet_11_::RAB-3 and CLA-1::7×GFP_11_ were reminiscent to those of transgenically expressed TdTomato::RAB-3 and CLA-1::3×GFPnovo2; we observed CLA-1::7×GFP_11_ puncta that are localized at the tip of the presynaptic varicosity labeled with 4×wrmScarlet_11_::RAB-3 (Figure 3B). We also observed similar localization pattern of 8×wrmScarlet_11_::CLA-1 (data not shown). To test if overexpression of GFP_1-10_ and wrmScarlet_1-10_ affects the localization patterns and fluorescent intensities of CLA-1::7×GFP_11_ and 4×wrmScarlet_11_::RAB-3, we compared the transgenic animals carrying *itr-1*p *(DA9 specific promoter)::wrmScarlet_1-10_::SL2::GFP_1-10_* extrachromosomal arrays generated at 20 ng/μL and 200 ng/μL concentrations. We did not observe a significant difference in the localization patterns and fluorescence intensity of CLA-1::7×GFP_11_ between animals injected at 20 ng/μL and 200 ng/μL concentrations (Figure S2 and Figure 3C). This is consistent with the idea that the endogenous expression level of CLA-1 defines the maximum signal intensity. For 4×wrmScarlet_11_::RAB-3, we did not observe a difference in the localization pattern, however, we noticed a tendency of higher fluorescence intensity of 4×wrmScarlet::RAB-3 in the animals injected at higher concentrations of *wrmScarlet_1-10_::SL2::GFP_1-10_* (Figure S2). This suggests that 4×wrmScarlet_11_::RAB-3 is not fully saturated with wrmScarlet_1-10_ in animals injected with *wrmScarlet_1-10_::SL2::GFP_1-10_* at 20 ng/μL and therefore have not yet reached the maximum fluorescence intensity set by the endogenous expression level of RAB-3. Nevertheless, our result suggests that overexpression of wrmScarlet_1-10_ and GFP_1-10_ does not affect localization of endogenously labeled synaptic proteins. We note a dim green signal in the cell body of DA9 of *mizIs41 (mig-13*p*::wrmScarlet1-10::SL2::GFP1-10)* animals (Figure 3D), consistent with previous observations that GFP_1-10_ by itself has a weak fluorescence^16^.

In DA9, presynaptic localization of CLA-1 and RAB-3 depends on *syd-2*^13^. Consistent with previous studies, the localization of TdTomato::RAB-3 and CLA-1::3×GFPnovo2 are greatly diminished from the axon in *syd-2(ok217)* mutants (Figure 3E). Similarly, signals of endogenously labeled CLA-1::7×GFP_11_ and 4×wrmScarlet_11_::RAB-3 were greatly diminished in the *syd-2* mutant; we observed dim diffuse signal of CLA-1::7×GFP_11_ and sporadic dim clusters of 4×wrmScarlet_11_::RAB-3 (Figure 3F). This further supports that the knock-in strains accurately recapitulate the localization patterns of CLA-1 and RAB-3, as described previously.

We used the *mizIs41 (mig-13*p*::wrmScarlet1-10::SL2::GFP1-10)* transgene to examine the localization patterns of other active zone proteins (SYD-2, UNC-10, and ELKS-1). The localization patterns of these active zone proteins tend to be inconsistent when labeling these proteins using traditional transgenic overexpression (unpublished observations). We found that the endogenously expressed 7×GFP_11_::SYD-2, UNC-10::7×GFP_11_, and ELKS-1::7×GFP_11_ were localized at the tip of synaptic varicosity labeled with 4×wrmScarlet::RAB-3 (Figures 3H-3J), similar to the CLA-1::7×GFP localization (Figure 3G). We also examined the co-localization between SYD-2 and CLA-1 in the *8×wrmScarlet_11_::syd-2; cla-1::7×GFP_11_; mizIs41* strain. We observed near-perfect co-localization between 8×wrmScarlet_11_::SYD-2 with CLA-1::7×GFP_11_ puncta (Figure 3K). This demonstrates that our endogenous labeling system is beneficial for labeling active zone proteins, which have been challenging with the traditional transgenic overexpression methods.

### Labeling presynaptic proteins in the D-type GABAergic motor neurons and PDE dopaminergic sensory neurons

We next examined if our knock-in strains can be used to visualize the presynaptic specializations in different neuron types. First, we examined the localization of endogenous CLA-1 and RAB-3 in the D-type GABAergic motor neurons (DDs and VDs). By the adult stage, DD and VD neurons have formed synapses onto the dorsal and ventral body wall muscles, respectively^45^ (Figure 4A). We expressed *wrmScarlet_1-10_::SL2::GFP_1-10_* under the GABAergic motor neurons specific promoter, *unc-25*p^46,47^ in *4×wrmScarlet_11_::rab-3; cla-1::7×GFP_11_* animals. Similar to DA9, we observed CLA-1::7×GFP_11_ puncta at the tip of presynaptic varicosity labeled with 4×wrmScarlet_11_::RAB-3 (Figures 4B and 4C). We next examined the localization patterns of endogenous CLA-1 and RAB-3 in the PDE dopaminergic sensory neurons by expressing *wrmScarlet_1-10_::SL2::GFP_1-10_* under the dopaminergic neuron-specific promoter, *dat-1*p^10,26^ in *4×wrmScarlet_11_::rab-3; cla-1::7×GFP_11_* animals. The PDE neurons are postembryonically-born bilaterally symmetric neurons, each of which extends an axon along the ventral nerve cord and forms *en passant* synapses onto the DVA interneuron^10,48^ (Figure 4D). Similar to previous observations using transgenically expressed RAB-3 and CLA-1^10^, the endogenously labeled 4×wrmScarlet_11_::RAB-3 and CLA-1::7×GFP_11_ puncta are colocalized along their axons (Figures 4E and 4F).

**Figure 4.**
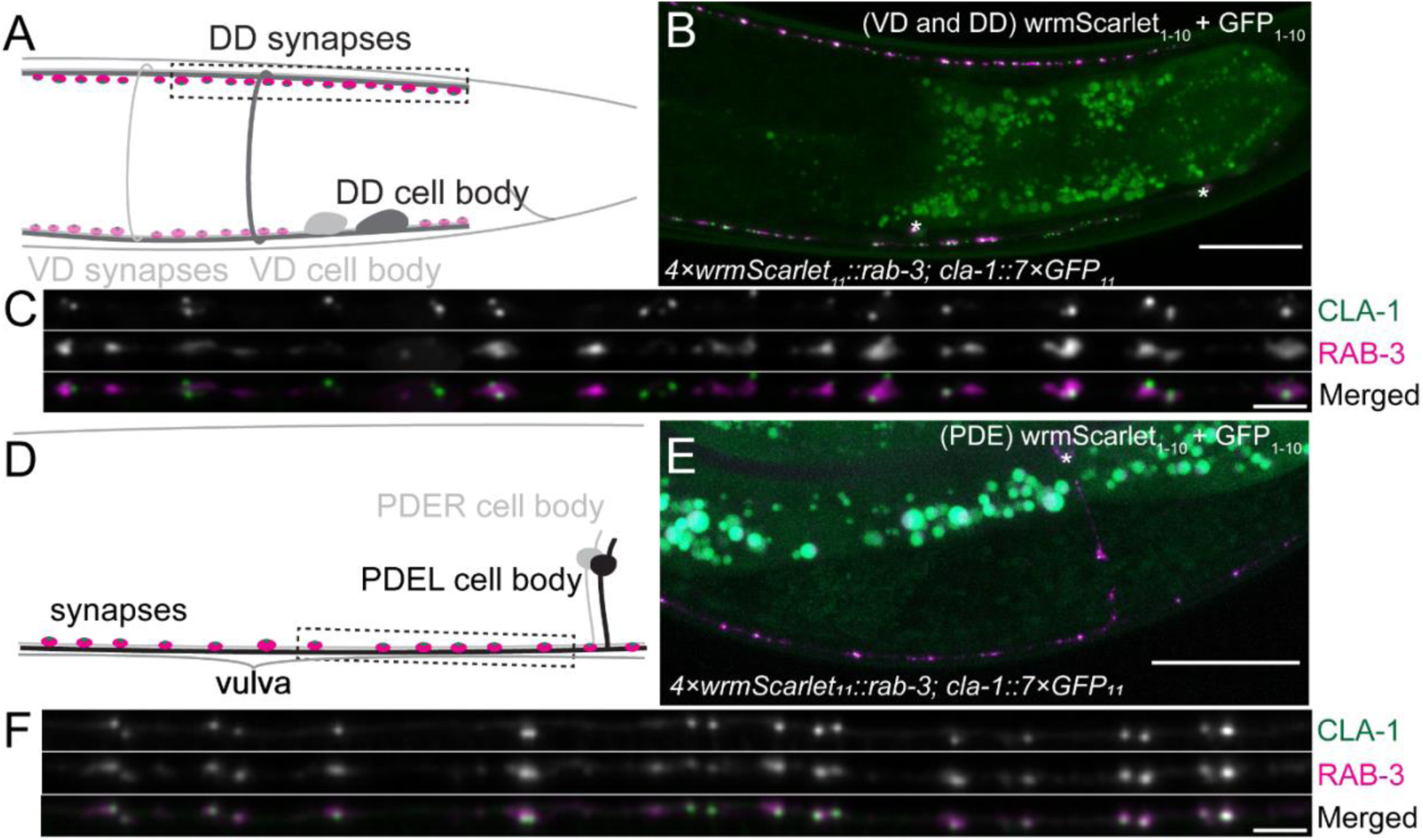
Labeling of endogenous CLA-1 and RAB-3 in DD/VD and PDE neurons. (A) The schematic of the most posterior VD (light grey) and DD (dark grey) motor neurons, VD13 and DD6. Magenta dots on the dorsal and ventral neurites represent presynaptic specializations of DDs and VDs, respectively. The dotted box represents the imaging area of the presynaptic domain of the DD6 neuron shown in (C.) (B) Representative image of the posterior VD and DD motor neurons labeled with endogenous 4×wrmScarlet::RAB-3 and CLA-1::7×GFP. The *unc-25* promoter was used to express GFP_1-10_ and wrmScarlet_1-10_. The asterisks denote VD and DD cell bodies. Scale bar: 10 μm. (C) Representative image of the DD6 presynaptic specializations labeled with endogenous CLA-1::7×GFP (top), 4×wrmScarlet::RAB-3 (middle), and merged image (bottom). Scale bar: 2 μm. (D) Schematic of the part of PDE neurons around the vulva. Magenta dots on the ventral neurites represent presynaptic specializations of the PDE neurons (PDEL and PDER). The dotted box represents the imaging area of the presynaptic domain of the PDE neuron shown in (F.) (E) Representative image of the part of the PDE neurons around vulva labeled with endogenous 4×wrmScarlet::RAB-3 and CLA-1::7×GFP. The *dat-1* promoter was used to express GFP_1-10_ and wrmScarlet_1-10_. The asterisk denotes the PDE cell body. Scale bar: 10 μm. (F) Representative image of the PDE presynaptic specializations labeled with endogenous CLA-1::7×GFP (top), 4×wrmScarlet::RAB-3 (middle), and merged image (bottom) in the axonal region anterior to the cell body and posterior to the vulva. Scale bar: 2 μm.

Together, we show that the localization pattern of endogenous 4×wrmScarlet_11_::RAB-3 and CLA-1::7×GFP_11_ are consistent among different neuron types, and the versatility of our system to label presynaptic proteins in different neuron types simply by using different promoters to express wrmScarlet_1-10_ and GFP_1-10_.

## Discussion

Here we generated a collection of strains in which sequences of tandem repeats of *GFP_11_* and/or *wrmScarlet_11_* are knocked into the loci of presynaptic genes. We show that transgenic expression of the remaining split fluorescent proteins from neuron- or neuron-type-specific promoters enables a robust and consistent visualization of the active zone proteins. We showed that overexpression of *wrmScarlet_1-10_* and *GFP_1-10_* does not affect the localization pattern and minimally affects the signal intensity of endogenously labeled presynaptic proteins. Comparisons between strains that transgenically overexpress synaptic proteins fused to fluorophores may be difficult as the fluorescence intensities and localization patterns are directly affected by differing expression levels of the transgene. In our strains, the maximal signal intensity of the labeled synaptic protein is determined by the endogenous expression level of the protein. Therefore, the use of these knock-in strains may provide a better standard for which labs can easily compare the qualitative differences of different genetic backgrounds in synapse formation and patterning. A CRISPR generated single copy insertion of *GFP_1-10_* and/or *wrmScarlet_1-10_* may further minimize the variability of the synaptic labeling, however, the ease of modularity of using different cell-specific promoters expressed using extrachromosomal arrays would be diminished.

Previous works have suggested that the mechanisms of presynaptic assembly vary among different neuron types^4^. For example, in the hermaphrodite-specific neurons (HSNs), the Arp2/3-dependent branched filamentous-actin (F-actin) functions as a structural scaffold to recruit active zone proteins including SYD-1 and SYD-2, which subsequently recruit other active zone proteins and SVs^49^. On the other hand, in the AIY interneurons, SYD-1 and SYD-2 are recruited to the synapses independent of F-actin ^50^. Our marker strains would be useful to exclude the possibility that the difference in presynaptic assembly in each neuron type is due to differential expression levels of the transgenes used to visualize active zone proteins.

### Limitations of the endogenous labeling system using split-fluorescent proteins

While our endogenous labeling platform provides robust visualization of the presynaptic proteins, there are several limitations to this labeling method as we discuss below.

First, our marker strains cannot distinguish/label certain isoforms of some active zone proteins. For example, CLA-1 has 6 isoforms, which are classified into 3 categories: CLA-1L (long), which includes CLA-1a and CLA-1b isoforms, CLA-1M (medium), which includes CLA-1c and CLA-1d isoforms, and CLA-1S (small) which includes CLA-1e and CLA-1f isoforms^13^. Transgenic labeling of CLA-1L and CLA-1S revealed a distinct subsynaptic localization of these isoforms^13,51^. Our knock-in strains labels CLA-1a, CLA-1d and CLA-1e isoforms, effectively labeling all 3 categories of CLA-1S/M/L isoforms. However, CLA-1b/c/f isoforms remain unlabeled, whose localization and functions have not been studied^13^. It is possible that these isoforms are expressed differently in different neurons which may account for neuron-type specific functions of CLA-1 in presynaptic assembly and functions^13^. It is therefore interesting to generate a series of knock-in strains to visualize specific isoforms of each active zone protein.

Second, a fusion of tandem repeats of GFP_11_ or wrmScarlet_11_ to endogenous presynaptic proteins could interfere with the protein functions. We show that endogenously tagged *unc-10, elks-1,* and *rab-3* strains exhibit mild to severe synaptic transmission defects. Extra caution should be paid when using these strains to examine synaptic functions in certain genetic backgrounds. Tagging *unc-10* and *elks-1* loci at their 5’ end may reduce the potential effects of GFP_11_ or wrmScarlet_11_ tags on their functions. For *rab-3* we note that our *4×wrmScarlet_11_::rab-3b* strain also labels *rab-3a* which is a relatively understudied isoform^40^. Although the expression of the *rab-3B* isoform is sufficient rescue *rab-3* mutants^40^, it is possible that *4×wrmScarlet_11_* inserted in the middle of *rab-3 A isoform* disrupted the functions of RAB-3 in neurotransmission. Alternatively, the functional defect of *4×wrmScarlet_11_::rab-3b* animals may be due to the *wrmScarlet_11_* insertion. Examination of GFP_11_ tagged *rab-3* would provide a clarification. C-terminal tagging of RAB-3 cannot be done as RAB-3 contains a CXC prenylation site required for proper association to the SV membrane^40,52^. Tagging other SV-associated proteins such as SNG-1/Synaptogyrin^53^, SNT/Synaptotagmin^54^ and SNN-1/Synapsin^55^ may provide a better labeling platform for visualizing SVs without functional perturbations.

## Data Availability

Strains and plasmids are available from the Caenorhabditis Genetics Center (CGC) and Addgene.

## Acknowledgements

We would like to thank Don Moerman for the general discussion and all of his work on the gene knockout consortium, including the generation of the *syd-2(ok217)* mutant strain, Peri Kurshan for providing the *wyIs685* transgenic strain, and Ardalan Hendi, Mo Miao, Sydney Ko, Menghao Lu for comments on the manuscript. We also thank the Mizumoto lab and colleagues for their general discussions. *syd-2(ok217)* used in this study were obtained from the *Caenorhabditis* Genetics Center, CGC, funded by the National Institute of Health (NIH) - Office of Research Infrastructure Programs (P40 OD010440) and *C. elegans* gene knockout consortium. This work is supported by the Canadian Institutes of Health Research (CIHR) and the Natural Sciences and Engineering Research Council of Canada (NSERC). M. K. is a recipient of CIHR CGS-D and the UBC 4-year fellowship. A. W. S. is a recipient of UBC Cell 1-year fellowship, and the Alzheimer Society Doctoral Award.

Supplemental Information

Genotyping primers:

*syd-2(ok217)*

F: CGCGGGAATTATGCCTATTA

R: AATCTCTAACCATGCGGTCG

Internal R: GCTTCTTCCGCTTCTGCTGC

*rab-3(miz237[4×wrmScarlet_11_::rab-3])*

F: ATCAGTTCCCTCTCGTTTCTC

R: CACCCTACCAACTAGGTCAAC

*cla-1(miz321[cla-1::7×GFP_11_]* and *miz378[cla-1::8×wrmScarlet_11_])*

F: CAGTGTCACTGGACATCGGCT

R: ACAACGGACCTACTACCCT

*syd-2(miz231[7×GFP_11_::syd-2] and miz329[8×mScarlet_11_::syd-2])*

F: ACAAGGGCAAGCTGATTCAC

R: TGTCCCGGTCTTCCAACATG

*elks-1(miz364[elks-1::7×GFP_11_]))*

F: TACCGGCTCCAGTGATTCC

R: TGTGTGCCATTGGATGTGAG

*unc-10(miz404[unc-10::7×GFP_11_])*

F: CGAACTCGGATCTCAACCAC

R: TTGGATGCACCGATTAGCTG

CRISPR:

*rab-3* gRNA: TCTGAAAATAGGGCTACTGT

*4×wrmScarlet11::rab-3* HDR template

GCTCTTTTAAAATAAATCTACAGTAGCCCTATTTTCAGATGTATACAGTTGTGGAACAATACGAGAAGTCCGTGGCCCGACATTGCACAGGCGGAGGTGGAAGTGGTGGCTACACGGTTGTAGAACAGTACGAGAAGAGTGTCGCACGTCATTGTACTGGCGGTGGAGGCTCTGGAGGATACACCGTAGTAGAGCAATATGAGAAAAGTGTGGCTCGTCATTGTACCGGCGGAGGAGGTTCCGGCGGATACACAGTGGTAGAACAGTACGAAAAGAGTGTCGCAAGACATTGTACAGGTGGCGCGGCTGGCGGACAACCTCAAGGCGCTACACCGGGACAAC

F:

GCTCTTTTAAAATAAATCTACAGTAGCCCTATTTTCAGATGTATACAGTTGTGGAACAATACG

R:

GTTGTCCCGGTGTAGCGCCTTGAGGTTGTCCGCCAGCCGCGCCACCTGTACAATGTCTTGCG

*cla-1* gRNA: GCAGTTTTCAGTTTATTGCA

*cla-1::7×GFP_11_* HDR template

GTCTTTTTTTTAAATATCTAAATCATTTAAATTTTTCAGTGTCACTGGACATCGGCTACCCTGCAATAAACGGTGCAGGAGCTGGAGCTGGAGCCGGAGCCGGAGCCCGTGACCACATGGTCCTTCATGAGTATGTAAATGCTGCTGGGATTACAGGTGGCTCTGGAGGTAGAGATCATATGGTTCTCCACGAATACGTTAACGCCGCAGGCATCACTGGCGGTAGTGGAGGACGCGACCATATGGTACTACATGAATATGTCAATGCAGCCGGAATAACCGGAGGGTCCGGAGGCCGGGATCACATGGTGCTGCATGAGTATGTGAACGCGGCGGGTATAACTGGTGGGTCGGGCGGACGAGACCATATGGTGCTTCACGAATACGTAAACGCAGCTGGCATTACTGGCGGATCAGGTGGCAGGGATCACATGGTACTCCATGAGTACGTGAACGCTGCTGGAATCACAGGCGGTAGCGGCGGTCGGGACCATATGGTCCTGCACGAATATGTCAATGCTGCCGGTATCACCGCAGCTGGAGGTTGAAAACTGCTCATAATGCTCAAAAATCTTTCTCAAAAGTTGACCAAAAAGCTCAAAAACTCAAACTTCT

F:

GTCTTTTTTTTAAATATCTAAATCATTTAAATTTTTCAGTGTCACTGGACATCGGCTACCCTGCAATAAACGGTGCAGGAGCTGGAGCTGGAG

R:

AGAAGTTTGAGTTTTTGAGCTTTTTGGTCAACTTTTGAGAAAGATTTTTGAGCATTATGAGCAGTTTTCAACCTCCAGCTGCGGTGATACCG

*cla-1::8×wrmScarlet_11_* HDR template

GTCTTTTTTTTAAATATCTAAATCATTTAAATTTTTCAGTGTCACTGGACATCGGCTATCCTGCAATAAACGGATCAGGATCTGGAAGTGGAAGCGGTGGATCTGGTGGATACACAGTTGTTGAGCAGTACGAGAAATCGGTTGCTCGACATTGCACAGGTGGTGGTGGAAGTGGTGGATATACTGTGGTGGAACAATACGAGAAGTCTGTGGCTAGACACTGTACTGGTGGTGGTGGTAGCGGAGGATACACAGTCGTCGAACAATACGAAAAGAGCGTTGCACGACACTGCACCGGTGGTGGTGGTTCTGGTGGTTATACAGTAGTAGAGCAATATGAGAAGAGTGTGGCTCGTCACTGCACTGGTGGTGGTGGTAGTGGTGGTTACACTGTTGTTGAACAATATGAGAAAAGCGTCGCCCGCCACTGTACAGGTGGTGGTGGATCCGGTGGATACACCGTAGTGGAACAGTATGAGAAATCAGTGGCCCGTCATTGCACCGGTGGTGGTGGATCTGGTGGATACACAGTGGTAGAACAATATGAGAAATCGGTGGCACGGCACTGTACTGGTGGTGGTGGAAGCGGTGGTTATACCGTCGTTGAGCAATACGAAAAATCAGTCGCCAGACACTGCACAGGTGGTTGAAAACTGCTCATAATGCTCAAAAATCTTTCTCAAAAGTTGACCAAAAAGCTCAAAAACTCAAACTTCT

F:

GTCTTTTTTTTAAATATCTAAATCATTTAAATTTTTCAGTGTCACTGGACATCGGCTATCCTGCAATAAACGGATCAGGATCTGGAAGTGGAAGCGG

R:

AGAAGTTTGAGTTTTTGAGCTTTTTGGTCAACTTTTGAGAAAGATTTTTGAGCATTATGAGCAGTTTTCAACCACCTGTGCAGTGTCTGGCGACT

*syd-2* gRNA: AGAAATATGAGCTACAGCAA

*7×GFP11::syd-2* HDR template

GAAATTTAGTTATCAATTTTAATCTGTTTCAGAAATATGGCCCGTGACCACATGGTCCTTCATGAGTATGTAAATGCTGCTGGGATTACAGGTGGCTCTGGAGGTAGAGATCATATGGTTCTCCACGAATACGTTAACGCCGCAGGCATCACTGGCGGTAGTGGAGGACGCGACCATATGGTACTACATGAATATGTCAATGCAGCCGGAATAACCGGAGGGTCCGGAGGCC GGGATCACATGGTGCTGCATGAGTATGTGAACGCGGCGGGTATAACTGGTGGGTCGGGCGGACGAGACCATATGGTGCTTCACGAATACGTAAACGCAGCTGGCATTACTGGCGGATCAGGTGGCAGGGATCACATGGTACTCCATGAGTACGTGAACGCTGCTGGAATCACAGGCGGTAGCGGCGGTCGGGACCATATGGTCCTGCACGAATATGTCAATGCTGCCGGTATCACCGGAGCCAGCTACAGCAATGGAAACATAAATTGTGATATAATGCCGACAATCT

F:

GAAATTTAGTTATCAATTTTAATCTGTTTCAGAAATATGGCCCGTGACCACATGGTCCTTCATGAGTATG

R:

GATTGTCGGCATTATATCACAATTTATGTTTCCATTGCTGTAGCTGGCTCCGGTGATACCGGCAGCATTGACATATTCG

*8×wrmScarlet11::syd-2* HDR template

AACATTTATTAATGTTTTGTTTTTGATTGAAAATCTGAAATTTAGTTATCAATTTTAATCTGTTTCAGAAATATGGGATCAGGATCTGGAAGTGGAAGCGGAGGATCTGGTGGATACACAGTTGTTGAGCAGTACGAGAAATCGGTTGCTCGACATTGCACAGGTGGTGGTGGAAGTGGTGGATATACTGTGGTGGAACAATACGAGAAGTCTGTGGCTAGACACTGTACTGGTGGTGGTGGTAGCGGAGGATACACAGTCGTCGAACAATACGAAAAGAGCGTTGCACGACACTGCACCGGTGGTGGTGGTTCTGGTGGTTATACAGTAGTAGAGCAATATGAGAAGAGTGTGGCTCGTCACTGCACTGGTGGTGGTGGTAGTGGTGGTTACACTGTTGTTGAACAATATGAGAAAAGCGTCGCCCGCCACTGTACAGGTGGTGGTGGATCCGGTGGATACACCGTAGTGGAACAGTATGAGAAATCAGTGGCCCGTCATTGCACCGGTGGTGGTGGATCTGGTGGATACACAGTGGTAGAACAATATGAGAAATCGGTGGCACGGCACTGTACTGGTGGTGGTGGAAGCGGTGGTTATACCGTCGTTGAGCAATACGAAAAATCAGTCGCCAGACACTGCACAGGTGGTGGATCATCTGGATCTGGATCCGGTAGCTATAGTAATGGAAACATAAATTGTGATATAATGCCGACAATCTCGGAAGATGGAGTGGACAACGGCGG TCCCA

F:

AACATTTATTAATGTTTTGTTTTTGATTGAAAATCTGAAATTTAGTTATCAATTTTAATCTGTTTCAGAAATATGGGATCAGGATCTGGAAGTGGAAGC

R:

TGGGACCGCCGTTGTCCACTCCATCTTCCGAGATTGTCGGCATTATATCACAATTTATGTTTCCATTACTATAGCTACCGGATCCAGATCCAGATGATCC

*elks-1* gRNA: TCGTCGTGATCTACCTGTGG

*elks-1::7×GFP_11_* HDR template

GTACCATTGGCATTCCACAACACTCTCAGCACCCGCCGCAAGTAGATCACGACGATGCTGACGGAATTTGGGCCGGTGCAGGAGCTGGAGCTGGAGCCGGAGCCGGAGCCCGTGACCACATGGTCCTTCATGAGTATGTAAATGCTGCTGGGATTACAGGTGGCTCTGGAGGTAGAGATCATATGGTTCTCCACGAATACGTTAACGCCGCAGGCATCACTGGCGGTAGTGGAGGACGCGACCATATGGTACTACATGAATATGTCAATGCAGCCGGAATAACCGGAGGGTCCGGAGGCCGGGATCACATGGTGCTGCATGAGTATGTGAACGCGGCGGGTATAACTGGTGGGTCGGGCGGACGAGACCATATGGTGCTTCACGAATACGTAAACGCAGCTGGCATTACTGGCGGATCAGGTGGCAGGGATCACATGGTACTCCATGAGTACGTGAACGCTGCTGGAATCACAGGCGGTAGCGGCGGTCGGGACCATATGGTCCTGCACGAATATGTCAATGCTGCCGGTATCACCGCAGCTGGAGGTTGAAAATTGTCTAGAATTGTCTGAACTTTTCCGAAATTTCCCAGTTTTTTCTCATATCTGTTCACTTACTTTTT

F:

GTACCATTGGCATTCCACAACACTCTCAGCACCCGCCGCAAGTAGATCACGACGATGCTGACGGAATTTGGGCCGGTGCAGGAGCTGGAGCTGGAGCCGG

R:

AAAAAGTAAGTGAACAGATATGAGAAAAAACTGGGAAATTTCGGAAAAGTTCAGACAATTCTAGACAATTTTCAACCTCCAGCTGCGGTGATACCGGCA

*unc-10* gRNA: TTTTTTGTCTTAATTACCAC

*unc-10::7×GFP_11_ HDR template*

AAGATTCCGATGTATCAGTTGGAGGTGCTCAGCAGGGAGCCCGTGACCACATGGTCCTTCATGAGTATGTAAATGCTGCTGGGATTACAGGTGGCTCTGGAGGTAGAGATCATATGGTTCTCCACGAATACGTTAACGCCGCAGGCATCACTGGCGGTAGTGGAGGACGCGACCATATGGTACTACATGAATATGTCAATGCAGCCGGAATAACCGGAGGGTCCGGAGGCCGGGATCACATGGTGCTGCATGAGTATGTGAACGCGGCGGGTATAACTGGTGGGTCGGGCGGACGAGACCATATGGTGCTTCACGAATACGTAAACGCAGCTGGCATTACTGGCGGATCAGGTGGCAGGGATCACATGGTACTCCATGAGTACGTGAACGCTGCTGGAATCACAGGCGGTAGCGGCGGTCGGGACCATATGGTCCTGCACGAATATGTCAATGCTGCCGGTATCACCTAAcaaatttcatatgtttttgtttgttttttgtcttaattaccacaCgtcatttctctctttctatcgtcattttctt

F:

AAGATTCCGATGTATCAGTTGGAGGTGCTCAGCAGGGAGCCCGTGACCACATGGTCC TTCA

R:

AAGAAAATGACGATAGAAAGAGAGAAATGACGTGTGGTAATTAAGACAAAAAACAAACAAAAACATATGAAATTTGTTAGGTGATACCGGCAGCATTGA

---

**Figure S1.**
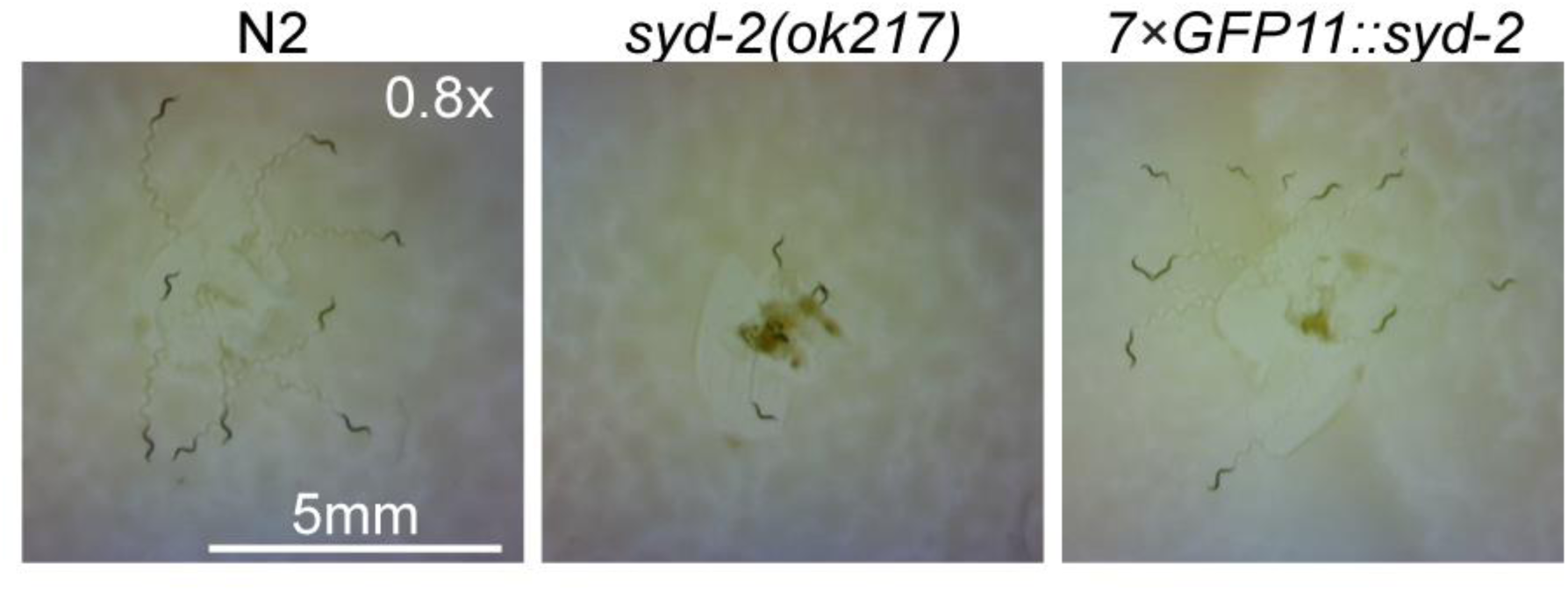
*7×GFP_11_::syd-2* animals exhibit normal locomotion. 9-10 wild type (N2), *syd-2(ok217), syd-2(miz231[7×GFP_11_::syd-2)* animals at the 4^th^ larval stage were placed onto the NGM plates, and images were taken 20 seconds after placing the animals.

**Figure S2.**
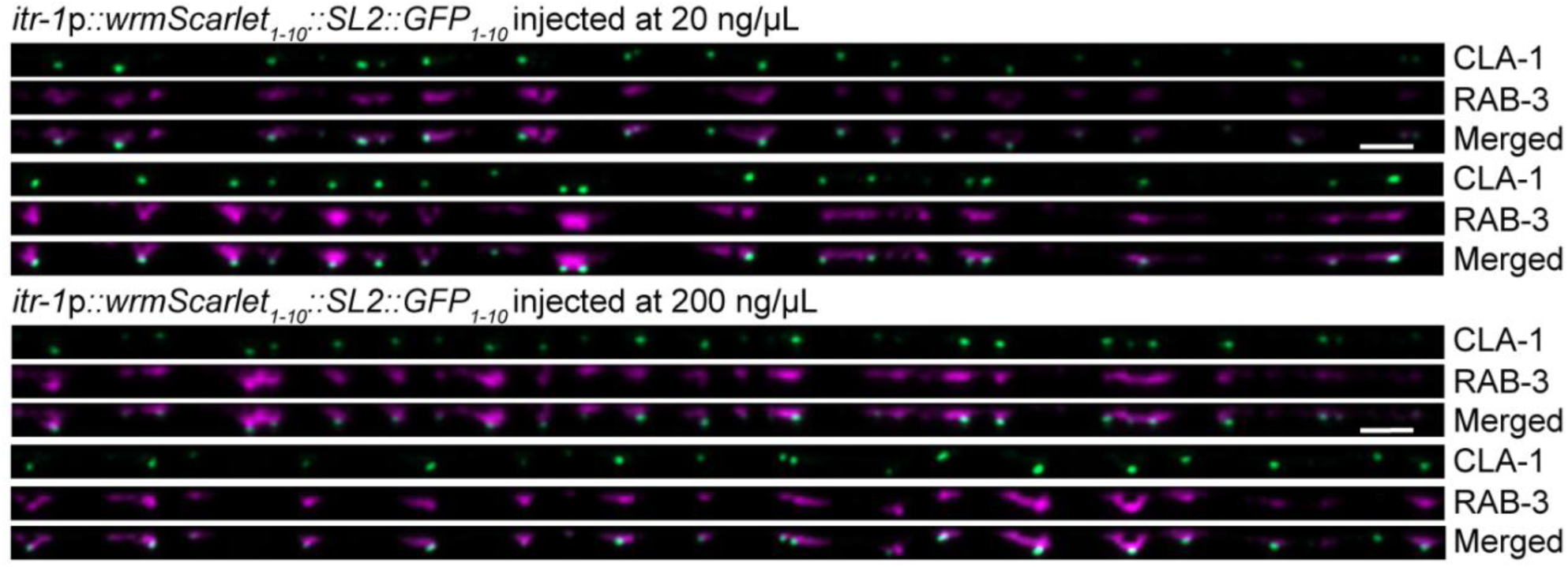
Overexpression of *wrmScarlet_1-10_::SL2::GFP_1-10_* does not affect the localization pattern of RAB-3 and CLA-1. Representative images of the DA9 presynaptic specializations labeled with endogenous 4×wrmScarlet::RAB-3 and CLA-1::7×GFP. *4×wrmScarlet::rab-3; cla-1::7×GFP* animals were injected with *itr-1*p::wrmScarlet_1-10_::SL2::GFP_1-10_ at 20 ng/uL (top) and 200 ng/uL (bottom). We noticed a tendency of 4×wrmScarlet::RAB-3 to be brighter in animals injected at higher concentrations. Scale bar: 2 um.

